# Screening autism-associated environmental factors in differentiating human neural progenitors with fractional factorial design-based transcriptomics

**DOI:** 10.1101/2022.06.27.497311

**Authors:** Abishek Arora, Martin Becker, Cátia Marques, Marika Oksanen, Danyang Li, Francesca Mastropasqua, Michelle Evelyn Watts, Manish Arora, Anna Falk, Carsten Oliver Daub, Ingela Lanekoff, Kristiina Tammimies

**Affiliations:** Center of Neurodevelopmental Disorders (KIND), Centre for Psychiatry Research, Department of Women’s and Children’s Health, Karolinska Institutet, and Child and Adolescent Psychiatry, Stockholm Health Care Services, Stockholm County Council, Stockholm, Sweden; Astrid Lindgren Children’s Hospital, Karolinska University Hospital, Region Stockholm, Stockholm, Sweden; Department of Chemistry - BMC, Uppsala University, Uppsala, Sweden; Department of Environmental Medicine and Public Health, Icahn School of Medicine at Mount Sinai, New York, USA; Department of Neuroscience, Karolinska Institutet, Stockholm, Sweden; Lund Stem Cell Center, Division of Neurobiology, Department of Experimental Medical Science, Lund University, Lund, Sweden; Department of Biosciences and Nutrition, Karolinska Institutet, Stockholm, Sweden; Science for Life Laboratory, Stockholm, Sweden

## Abstract

Research continues to identify genetic variation, environmental exposures, and their mixtures underlying different diseases and conditions. There is a need for screening methods to understand the molecular outcomes of such factors. Here, we investigate a highly efficient and multiplexable, fractional factorial experimental design (FFED) to study six environmental factors and four human induced pluripotent stem cell line derived differentiating human neural progenitors. We showcase the FFED coupled with RNA-sequencing to identify the effects of low-grade exposures to these environmental factors and analyse the results in the context of autism spectrum disorder (ASD). We performed this after five-day exposures on differentiating human neural progenitors accompanied by a layered analytical approach and detected several convergent and divergent, gene and pathway level responses. We revealed significant upregulation of pathways related to synaptic function and lipid metabolism following lead and fluoxetine exposure, respectively. The lipid changes were validated using mass spectrometry-based metabolomics after fluoxetine exposure. Our study demonstrates that the FFED can be used for multiplexed transcriptomic analyses to detect relevant pathway-level changes in human neural development caused by low-grade environmental risk factors. Future studies will require multiple cell lines with different genetic backgrounds for characterising the effects of environmental exposures in ASD.

## Introduction

Intensified research continues to identify genetic variation, environmental exposures and their mixtures underlying different diseases and conditions. As several factors can be associated with the same outcome, there is a need for better methods to investigate the molecular, cellular, and developmental effects of these factors, both in parallel and as mixtures. At present, there is a growing need to develop *in-vitro* experimental models of environmental factors that recapitulate real-life exposures and clinical outcomes (Austin, 2019; Caporale et al., 2022; Curtin et al., 2018).

Fractional factorial experimental designs (FFED) (Mee, 2009), adapted from methods used in managerial and industrial processes, drastically reduce the number of samples required to observe effects with high statistical power. It is a subset of a full factorial design that increases experimental efficiency and makes it possible to study the interactions between different experimental conditions using statistical modelling. Earlier, it has not been coupled with omic readouts such as RNA sequencing to investigate any changes in the transcriptional landscape.

Autism spectrum disorder (ASD) is a neurodevelopmental disorder (NDD) diagnosed in nearly 1-2% of the population (Lord et al., 2020). More than 100 genes are associated with ASD affected by both, common variants with small effect sizes and rare variants with larger effect sizes (Iakoucheva et al., 2019; Lord et al., 2020; Satterstrom et al., 2020). Heritability studies show that genetic factors account for 50 to 86.8% of ASD risk, suggesting that the remaining risk arises from environmental factors (Bai et al., 2019). Early exposure to chemical stressors may induce or exacerbate neurodevelopmental trajectories underlying ASD. Environmental factors associated with ASD include lead (Pb) (Heyer and Meredith, 2017), valproic acid (VPA) (Christensen et al., 2013; Moore, 2000), bisphenol A (BPA) (Thongkorn et al., 2019, 2021), ethanol (EtOH) (Stevens et al., 2013), fluoxetine hydrochloride (FH) (Boukhris et al., 2016) and zinc dysregulation (Arora and Austin, 2013; Arora et al., 2017). The majority of genes and environmental factors associated with ASD have also been implicated in other NDDs, suggesting more generalised effects on neurodevelopment rather than being ASD specific.

Despite the growing interest in environmental factors, the molecular mechanisms leading to the increased liability of behavioural and cognitive dysfunctions remain largely unknown, especially in genetically vulnerable conditions. The use of induced pluripotent stem cell (iPSC) derived neural progenitors, either from typically developed individuals or individuals with ASD, can enable the study of the effects of environmental factors during neurodevelopment (Pintacuda et al., 2021). As a large number of both genetic and environmental factors have been indicated in ASD, there is a need for efficient methods to estimate their effects during early development.

Here, we present a multiplexed experimental design based on the FFED coupled with RNA-sequencing to perform simultaneous analyses of six environmental factors associated with ASD and four cell lines. The main outcomes were determined using transcriptional and pathway level analysis in differentiating neural progenitors from human iPSCs. Furthermore, we validated the specific lipid pathways identified after exposure to FH using mass spectrometry-based metabolome detection. We show that FFED-RNA-seq can help pinpoint relevant mechanisms related to ASD-associated environmental exposures, and that there is a need to model these effects in multiple genetic backgrounds.

## RESULTS

### Cellular effects of environmental factors

We evaluated the molecular effects of Pb, VPA, BPA, EtOH, FH and zinc deficiency (Zn-) in cell lines with different clinical backgrounds for ASD during differentiation of human neural progenitors (Figure 1A). The Non-ASD cell lines included two neurotypical controls (CTRL_Male_, CTRL_Female_), while the ASD cell lines included two males with a known ASD genetic variant (ASD_CASK_, ASD_HNRNPU_) as described earlier (Becker et al., 2020; Falk et al., 2012; Mastropasqua et al., 2022; Uhlin et al., 2017). For studying the effects of the six exposures, we set up an FFED framework (Table S1A).

**Figure 1.**
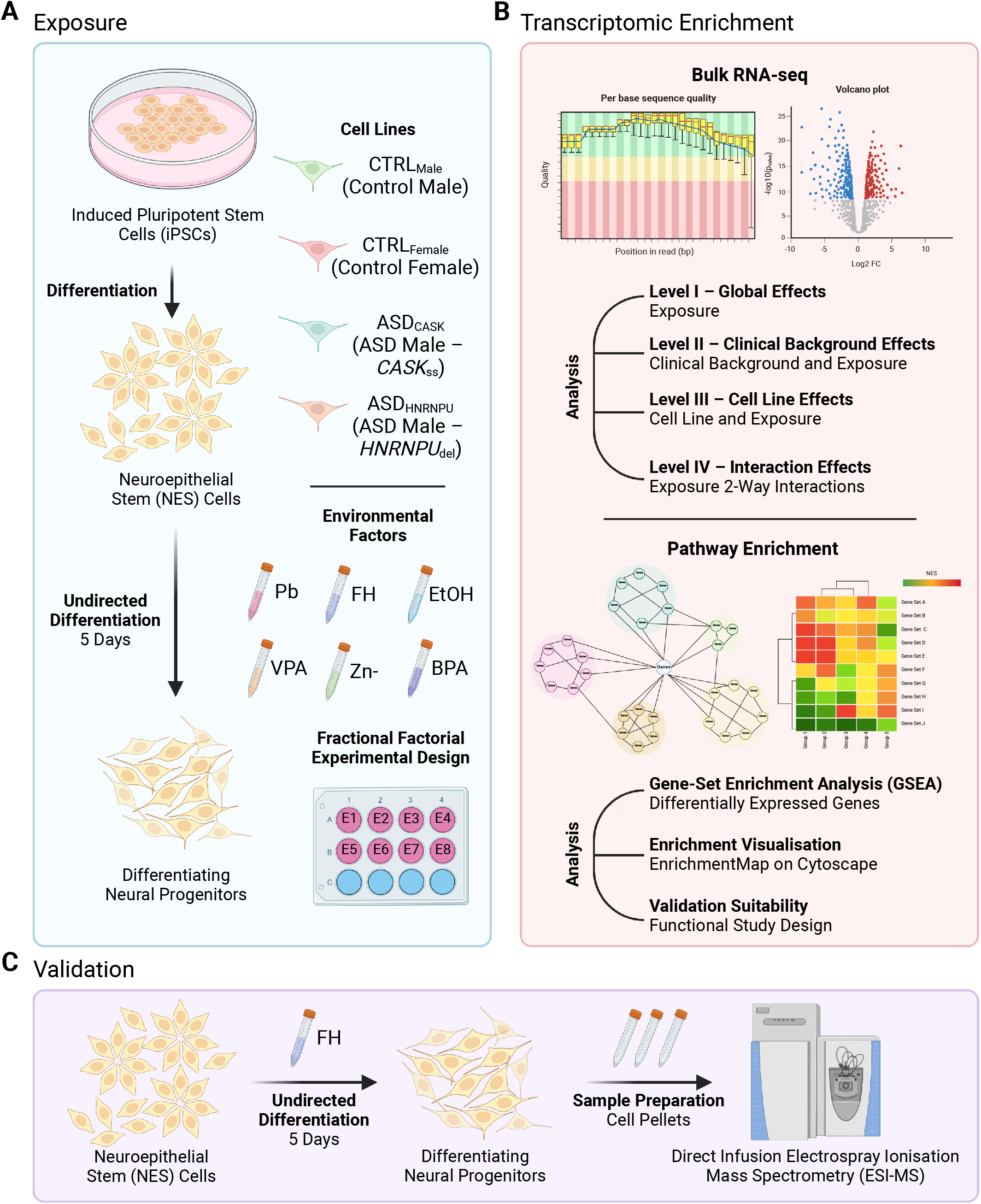
Overview of study plan using fractional factorial experimental design (FFED) (A) Exposure of neuroepithelial stem (NES) cells derived from human induced pluripotent stem cells (iPSCs) during neural progenitor differentiation for 5 days. Four cell lines, neurotypical controls (CTRL_Male_, CTRL_Female_) and males with autism spectrum disorder (ASD) diagnoses (ASD_CASK_: *CASK* splice site variant, ASD_HNRNPU_: *HNRNPU* deletion), were exposed to six environmental factors during differentiation, namely lead (Pb), fluoxetine hydrochloride (FH), ethanol (EtOH), valproic acid (VPA), bisphenol A (BPA) and zinc deficiency (Zn-). (B) Using RNA samples from (A), bulk RNA-sequencing was performed followed by differential gene expression analysis for global effects, clinical background effects, cell line effects and interaction effects. Using this, pathway enrichment and network visualisation was done. (C) Enriched pathways identified from (B) following FH exposure were detected using direct infusion electrospray ionisation mass spectrometry (ESI–MS) and quantified across the different levels of analyses.

First, we performed a cytotoxicity assay to test the exposure ranges of the six selected environmental factors in differentiating human iPSC-derived neuroepithelial stem (NES) cells. NES cells are neural progenitors that are predominantly neurogenic in nature but also have the potential to be gliogenic, under defined differentiation conditions (Chambers et al., 2009; Falk et al., 2012). Exposure ranges were selected based on an extensive literature review as described in Table S1B–S1F and tested in the CTRL_Male_ cell line. The highest concentration with no significant (Tukey post hoc p>0.05) or trending decrease in cell viability and no gross changes in cell morphology at 120 hours of exposure was selected for each environmental factor (Figure S1A); 3 µM for Pb, BPA and FH, and 3 mM for VPA and EtOH. This was done to avoid the toxicity of a single environmental factor overshadowing the combined effects of the other factors. Based on the six factors, we fitted the L8 Orthogonal Array based FFED for our study (Table S1A). The selected concentrations were then tested for cytotoxicity using the MTS assay in the FFED. At 120 hours of exposure, a significant difference (Tukey post hoc p<0.05) in cell viability was only observed for Pb, VPA and EtOH (Figure S1B). Based on the observed cytotoxicity effects of VPA on gross cellular morphology during NES cell differentiation in the FFED, the exposure concentration was adjusted to 3.0 µM in line with previous *in-vitro* exposures reported in the literature (Table S1D).

Next, we tested if the exposures affected the population of neural progenitors in two of the four cell lines, CTRL_Male_ and ASD_CASK_, by analysing the percentage of SOX2 positive cells at day 5 of exposure using the FFED. No significant changes (Tukey post hoc p>0.05) in the number of SOX2 positive cells were detected for any of the exposures (Figure S1C). Additionally, we tested for any proliferation changes using the BrdU assay and FFED at day 5 of exposure in all the cell lines. A significant decrease in proliferation was detected after exposure to Pb in the ASD_CASK_ cell line (Tukey post hoc p<0.001, Figure S1D).

### Approaching multiplexed exposures and gene expression

We investigated if FFED coupled with RNA-seq could detect relevant transcriptomic changes after low-grade environmental exposures on differentiating human neural progenitors for 5 days. We performed RNA-seq using the L8 orthogonal array for the four cell lines (n=32). We included a biological replicate of the full set for CTRL_Male_ (n=8) and a partial set for CTRL_Female_ (n=4), for variability and sample size testing. Principal component analysis (PCA) of the RNA-seq data (N=43) showed that the genetic background of the samples was the main driver of the differences across PC1 (46.12%) and PC2 (33.57%), and the exposure effects were minimal for the total variation in the transcriptomic profiles (Figure S2A and S2B).

We then performed identification of differentially expressed genes (DEGs) and gene-set enrichment analysis (GSEA) for enriched pathways at three levels. Briefly, at *Level I – Global Effects*, exposure effects were analysed across all cell lines; *Level – II, Clinical Background Effects*, exposure comparisons were made separately for two groups: non-ASD and ASD cell lines; and *Level III – Cell Line Effects*, exposure effects were analysed for every individual cell line (Figure 1B).

### Early exposure effects of environmental factors

In the global analyses (*Level I)*, no genes responding to BPA, VPA, EtOH and Zn-exposures (adj. p>0.05) were identified. Nevertheless, pathway analysis highlighted biological processes following these exposures (Table S3A and S3B). For BPA, twelve pathways were significantly upregulated, including protein lipid complex assembly (q=0.0027) and mitochondrial membrane organisation (q=0.032), and one pathway was significantly downregulated, detection of chemical stimulation involved in sensory perception (q=0.035). For VPA, thirty pathways were significantly downregulated, including neuron projection guidance (q=0.0023) and axon development (q=0.0089). For EtOH and Zn-, negative regulation of endothelial cell apoptotic process (q=0.0017) and positive chemotaxis (q=0.048) were the only significantly upregulated pathways, respectively. Additionally, for Zn-three pathways were significantly downregulated: plasminogen activation (q=0.0006), protein activation cascade (q=0.0048) and fibrinolysis (q=0.0066).

We detected significant changes in genes responding to Pb (69 genes, adj. p<0.05, Table S2A, Figure 2A) and FH (50 genes, adj. p<0.05, Table S2H, Figure 2C). Furthermore, pathway analysis revealed several biological processes affected by the exposures (Figure 2B and 2D). Amongst the upregulated pathways after Pb exposure (Table S3A) were cholinergic synaptic transmission (q=0.0099), axon extension (q=0.012), and synapse assembly (q= 0.012). Significantly downregulated pathways (Table S3B) were negative regulation of receptor signalling pathway via STAT (q=0.0091), regulation of heterotypic cell adhesion (q= 0.043) and axoneme assembly (q=0.046). For FH (Table S3A), the significantly upregulated pathways included alcohol biosynthetic process (q<0.0001), sterol biosynthetic process (q<0.0001), and regulation of steroid metabolic process (q<0.0001). There were six significantly downregulated pathways detected (Table S3B), including motile cilium assembly (q=0.015), axonemal dynein complex assembly (q=0.018) and cilium movement (q=0.035).

**Figure 2.**
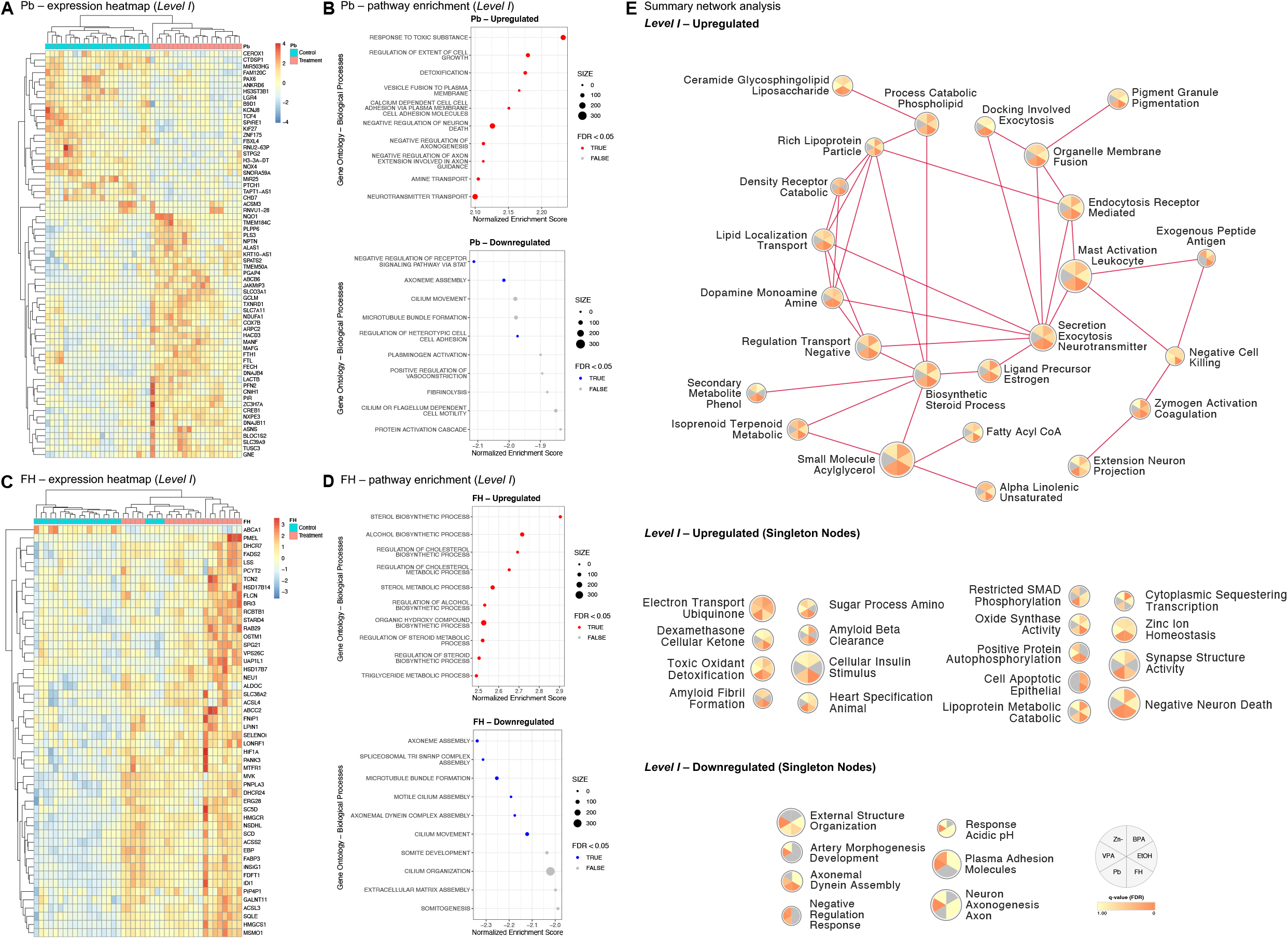
Differential gene expression and pathway enrichment – *Level I* analysis of global effects following environmental factor exposure. (A) Heatmap of significant (adj. p<0.05) differentially expressed genes following Pb exposure for 5 days. (B) Gene Ontology (GO) enrichment analysis with top 10 upregulated and downregulated biological processes following Pb exposure for 5 days. (C) Heatmap of significant (adj. p<0.05) differentially expressed genes following FH exposure for 5 days. (D) GO enrichment analysis with top 10 upregulated and downregulated biological processes following FH exposure for 5 days. (E) Summary network of upregulated and downregulated clusters enriched for day 5 exposure of lead (Pb), fluoxetine hydrochloride (FH), ethanol (EtOH), valproic acid (VPA), bisphenol A (BPA) and zinc deficiency (Zn-).

To assess the involvement of the genes responding to Pb and FH exposures across spatio-temporal trajectories during human neurodevelopment, we analysed their expression using available RNA-seq data from the BrainSpan Consortium (Miller et al., 2014). The genes responding to Pb (Figure S2C) and FH (Figure S2E) both formed two groups for early and late neurodevelopment. Both Pb (Figure S2D) and FH (Figure S2F) gene groups were significantly (adj. p<0.05) detected in the frontal cortex, temporo-parietal cortex, sensorimotor cortex, and subcortical regions. When considering their role across developmental periods, Pb genes (early) were significantly identified (adj. p<0.05) in all brain regions from the late mid-foetal period (19-24 weeks), other than in the subcortical region that was only significant (adj. p<0.05) from the early foetal period (0-12 weeks). For FH (late), the genes were significantly found (adj. p<0.05) in all studied brain regions, however, were limited to postnatal periods of neurodevelopment.

Additionally, we tested for enrichment of the responding genes in neurodevelopmental disorder gene lists, including SFARI for ASD (Banerjee-Basu and Packer, 2010), Genetic epilepsy syndromes (EPI) panel (Stark et al., 2021), intellectual disability (ID) panel (Stark et al., 2021) and a general developmental gene list (Li et al., 2021). No significant findings were observed (adj. p>0.05).

### Convergence of affected molecular pathways for all exposures

Several shared biological processes were revealed, when we analysed the molecular pathways across environmental factor exposures (*Level I* analysis). There was significant upregulation detected (Table S3A) in glycosphingolipid metabolic processes (FH q=0.034, Pb q=0.023), steroid catabolic processes (FH q=0.0003, Pb q=0.016), terpenoid metabolic processes (BPA q=0.046, FH q=0.031), negative regulation of endothelial cell apoptosis (EtOH q=0.0017, FH q=0.003), synaptic vesicle priming (FH q=0.014, Pb q=0.018) and cholinergic synaptic transmission (FH q=0.015, Pb q=0.01). Significantly downregulated biological processes (Table S3B) detected were axoneme assembly (FH q=0.0008, Pb q=0.046) and microtubule bundle formation (FH q=0.0065, Pb q=0.05). The summary network of the six exposures, indicated connected nodes across the exposures for several of the upregulated clusters and only singleton nodes for the downregulation (Figure 2E, Table S4A and S4B).

### Clinical background modulates exposure effects

Similar pathway analyses were performed for *Level – II, Clinical Background Effects*, by analysing Non-ASD cells (CTRL_Male_ and CTRL_Female_; Table S2B and S2I) and ASD cells (ASD_CASK_ and ASD_HNRNPU_; Table S2C and S2J) separately. The environmental factor exposures uniquely and commonly modulated several pathways. A total of 460 biological processes were upregulated (Table S3C) and 79 were downregulated (Table S3D) in the Non-ASD cells. In the ASD cells, 861 pathways were upregulated (Table S3E), while 17 pathways were downregulated (Table S3F). The summary networks were used to visualize hub nodes (Table S4C and S4D, Figure S3A and S3C for the Non-ASD group; Table S4E and S4F, Figure S3F and S3D for the ASD group).

Furthermore, we analysed the global network properties of the Non-ASD and ASD cell responses to the environmental factor exposures to represent the differential responses. In the upregulation network for Pb, the diameter measuring the network size, was greater for the ASD group (17.0) than the Non-ASD group (7.0). In contrast, the clustering coefficient indicating the nodal neighbourhood connectivity was higher of the Non-ASD group (0.68) than the ASD group (0.54). The average degree, representing the number of edges per node in a network, was conversely higher in the ASD group (8.06) than the Non-ASD group (6.67). Interestingly, no downregulation network could be generated for the ASD group based the selected thresholds (q<0.05). For the upregulation network of FH, the diameter of the Non-ASD group (18.0) was larger than the ASD group (4.0). The clustering coefficient was greater for the ASD group (0.95) than for the Non-ASD group (0.59), while the average degree was higher for the Non-ASD group (7.53) when compared to the ASD group (5.72). Similar to Pb, no downregulation network could be generated for FH exposure in the ASD group.

Lastly, we performed the *Level III* analyses for each cell line (Table S2D–S2G and S2K–S2N). Several pathways were found to be significantly modulated (Table S5). Interestingly, the exposures per cell line modulated several unique biological processes. For Pb, significantly (q<0.05) upregulated unique pathways (Table S5A), were indicated for all cell lines, however ASD_HNRNPU_ showed the largest effects with 1105 significantly upregulated pathways overall. While Pb-driven significantly downregulated pathways (Table S5B) had the largest effects in CTRL_Male_ with 266 out of 290 pathways that were significantly downregulated overall. In the case of FH, significantly (q<0.05) upregulated pathways (Table S5C) were mostly detected in CTRL_Male_ (250 out of 378 pathways) and none in ASD_CASK_. FH-driven significant downregulated pathways (Table S5D) were identified mostly in CTRL_Male_ (75 out of 85 pathways) with a few in CTRL_Female_, such as spliceosomal snRNP assembly (CTRL_Male_ q<0.0001) and cilium movement (CTRL_Female_ q=0.0002). Pb and FH exposure effects on the cell lines were visualised using summary network analysis for upregulated (Figure 3A and 3C, Table S4G and S4I, respectively) and downregulated (Figure 3B and 3D, Table S4H and S4J, respectively) common clusters.

**Figure 3.**
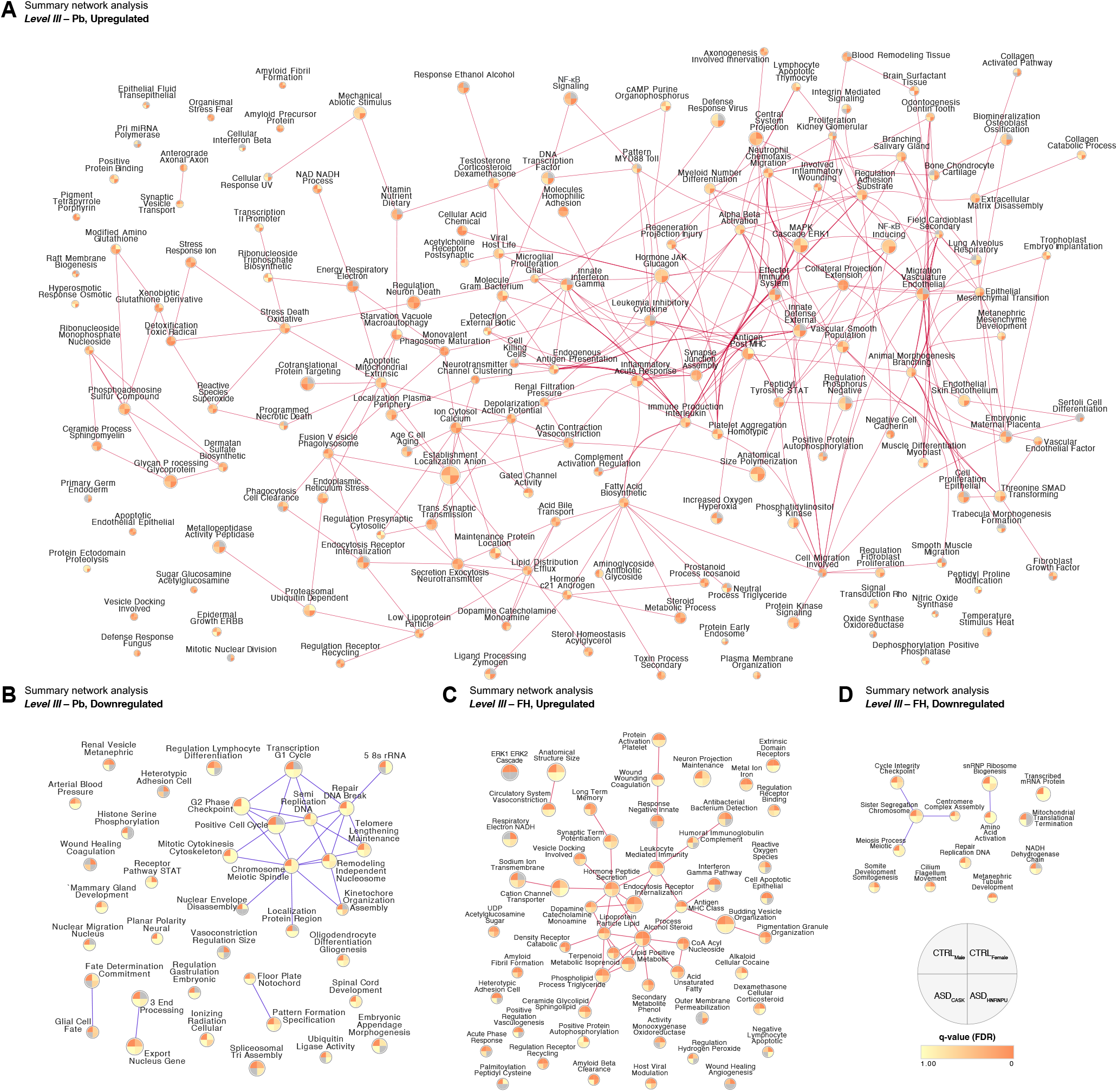
Summary networks from pathway enrichment – *Level III* analysis of cell line effects following lead (Pb) and fluoxetine (FH) exposure. (A) Upregulated and (B) downregulated clusters enriched for day 5 exposure of Pb in CTRL_Male_, CTRL_Female_, ASD_HNRNPU_ and ASD_CASK_. (C) Upregulated and (D) downregulated clusters enriched for day 5 exposure of FH in CTRL_Male_, CTRL_Female_, ASD_HNRNPU_ and ASD_CASK_.

BPA predominantly modulated pathways with significance (q<0.05) in CTRL_Male_ (upregulated 138 out of 139 pathways, downregulated 1 out of 1 pathway) (Table S5E). On the other hand, EtOH and VPA predominantly upregulated pathways (Table S5I and S5G, respectively) with significance in CTRL_Female_ (EtOH: 67 out of 72 pathways, VPA: 5 out of 5 pathways). VPA-driven significant downregulation (Table S5H) of pathways was detected mainly in CTRL_Female_ (37 out of 84 pathways) and ASD_CASK_ (51 out of 84 pathways). Zn-significantly upregulated pathways (Table S5J) mainly in CTRL_Male_ (21 out of 34 pathways), such as ribosome biogenesis (CTRL_Male_ q=0.0133) and translation initiation (CTRL_Male_ q=0.0332). The significantly downregulated pathways (Table S5K) like protein activation cascade (CTRL_Female_ q<0.0001) and sterol import (CTRL_Female_ q=0.0248) were primarily detected in CTRL_Female_ (34 out of 40 pathways).

### Detecting interaction effects with FFED

In addition to single exposures, the FFED design enabled exposure interaction analyses. In our design, we were able to analyse three two-way interactions (*Level IV* analyses): Pb – Zn-, VPA – FH, and BPA – EtOH. However, interaction modulated genes were only detected in CTRL_Male_ when analysing for interaction effects. This could be attributed to the presence of two full biological replicates of the FFED design for CTRL_Male_ (Table S1A). Several biological processes were modulated by the two-way interactions (Table S6A and S6B) and in the summary network several clusters were identified. Significant upregulation (Table S6C, Figure S3B) was detected in nucleosome organization assembly (BPA – EtOH q=0.035), mitochondrial electron respiration (Pb – Zn-q=0.015), membrane raft organization (Pb – Zn-q=0.040), lipoprotein density particle (VPA – FH q=0.023) and steroid biosynthetic process (VPA – FH q=0.034). Significant downregulation (Table S6D, Figure S3E) was detected in clusters such as G1 phase cell cycle (Pb – Zn-q=0.015), chromatin remodelling (Pb – Zn-q=0.0023), kinetochore organization assembly (Pb – Zn-q=0.0034, VPA – FH q=0.015), and snRNP biogenesis (VPA – FH q=0.0071).

### Lead and Bisphenol A induce alternative splicing events

After investigating gene response differences, we performed differential exon usage (DEU) analysis on the *Level I* to *III* comparisons (Table S7A). In line with the gene response analysis, Pb exposure presented significant effects on alternative splicing. Pb exposure induced more alternative splicing events in the ASD group (45 genes) compared to the Non-ASD (4 genes). In addition to Pb, BPA also affected alternative splicing. BPA exposure caused extensive alternative splicing events on the global level (223 genes) and in the Non-ASD group (346 genes). Furthermore, in the female control cell line, CTRL_Female_, alternative splicing events were detected following analysis in both the full set (381 genes) and excluded partial set (868 genes). No alternative splicing events were detected after exposure to the other environmental factors.

The genes with alternative splicing events in CTRL_Female_ after BPA exposure (partial set, 868 genes) were over-represented in several biological processes (Table S7B, Figure S3G). These included cytoskeleton organisation (adj. p=0.0091), neuron development (adj. p=0.036), regulation of RNA splicing (adj. p=0.0359), neuron differentiation (adj. p=0.048) and intracellular protein transport (adj. p=0.044). No significant enrichment (adj. p<0.05) in biological processes were detected for the genes driving alternative splicing events following Pb exposure at all levels of analyses.

Furthermore, genes with significant alternative splicing events following BPA exposure in CTRL_Female_ were tested for enrichment in publicly available gene lists, as previously done for significantly responding genes. Significant enrichment was observed in the SFARI gene list (adj. p=0.045), SFARI high confidence gene list (adj. p=0.022), ID panel (adj. p=0.0045) and developmental gene list (adj. p=0.0013). No significant enrichment (adj. p>0.05) was detected in the EPI panel.

### Fluoxetine alters lipid metabolism during neural progenitor differentiation

As we revealed an extensive dysregulation of lipid metabolism related pathways in the transcriptomic profiles after FH exposures, we set out to validate the findings using direct infusion electrospray ionisation mass spectrometry (ESI-MS) for metabolomics (Figure 1C). We initially evaluated over 2500 metabolite signals, including adducts, for FH exposure effects in all the cell lines, out of which 79 were selected and reported based on their signal threshold level. At the *Level I* analysis for global effects, we could confirm several lipid-based metabolites to be significantly elevated after exposure to FH, including fatty acids (FA 18:1, adj. p=0.0015; FA 20:2, adj. p=0.0056; FA 20:3, adj. p=0.0040), plasmanylcholines (PC e 32:0, adj. p=0.0006; PC e 32:1, adj. p=0.0003) and the phosphatidylethanolamine, PE 36:4 (adj. p=0.0062) (Figure 4A, Table S8A).

**Figure 4.**
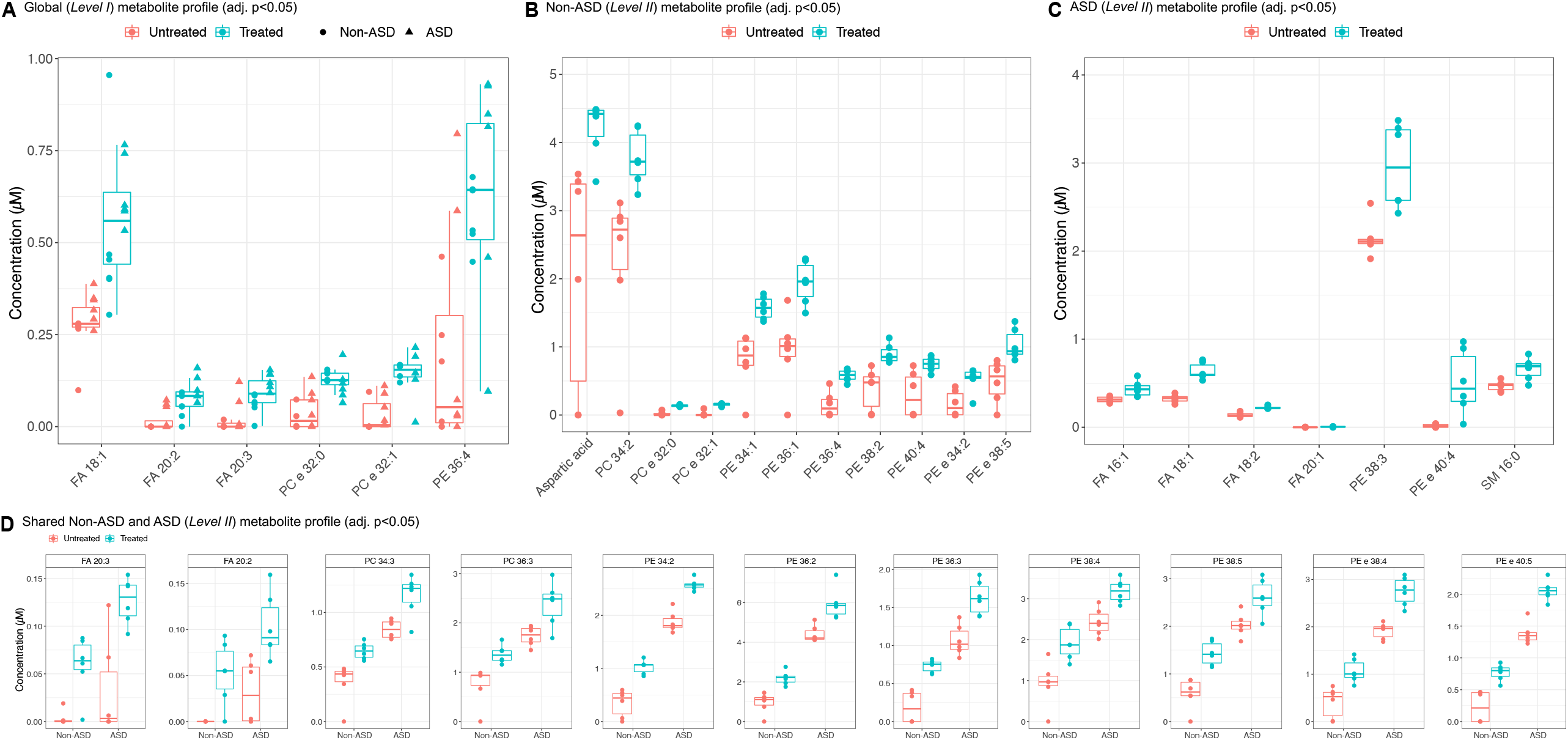
Metabolic changes induced by fluoxetine (FH) exposure. (A) Significantly enriched metabolites (adj. p<0.05) from day 5 FH exposure for global effects (*Level I* analysis). (B) Significantly enriched (adj. p<0.05) and unique metabolites from day 5 FH exposure, in the non-ASD (*Level II* analysis). (C) Significantly enriched (adj. p<0.05) and unique metabolites from day 5 FH exposure, in the ASD group (*Level II* analysis). (D) Significantly enriched (adj. p<0.05) and common metabolites from day 5 FH exposure, in the non-ASD and ASD groups (*Level II* analysis).

When separating for the clinical background (*Level* II) effects, several shared metabolites showed a significant elevation following FH exposure, however, with a difference in magnitude of concentrations, including fatty acids FA 20:2 (Non-ASD q=0.0217; ASD q=0.0320), FA 20:3 (Non-ASD q=0.0134, ASD q=0.0198); phosphatidylcholines PC 34:3 (Non-ASD q=0.0245, ASD q=0.0313), PC 36:3 (Non-ASD q=0.0246, ASD q=0.0432) and phosphatidylethanolamines PE 34:2 (Non-ASD q=0.0032, ASD q=0.0007), and plasmanylethanolamine PE e 38:4 (non-ASD q=0.0189, ASD q=0.0022) (Figure 4D, Table S8B and S8C).

Furthermore, we detected several metabolites that were elevated significantly only in either of the clinical background groups. For the Non-ASD cell lines, additional 11 metabolites showed significant changes, including aspartic acid (q=0.0436), PC e 32:0 (adj. p=0.0003), PE 34:2 (adj. p=0.0033) and PE e 40:5 (adj. p=0.0072) (Figure 4B, Table S8B). For the ASD cell lines, seven unique metabolites showed significant changes, including FA 18:2 (adj. p=0.0009), PE 38:3 (adj. p=0.0280), PE e 40:4 (adj. p=0.0426) and SM 16:0 (adj. p=0.0409) (Figure 4C, Table S8C).

At the *Level III* analysis for cell line effects, in addition to some of the earlier significant metabolites, few more metabolites were also significantly elevated. In CTRL_Male_ (Table S8D), PC e 32:0 (adj. p=0.020) and PC e 32.1 (adj. p=0.0068) were elevated upon FH exposure, while no metabolites were significantly elevated in CTRL_Female_ (Table S8E). In ASD_HNRNPU_ (Table S8F), significant elevation was detected in six metabolites including FA 20:3 (adj. p=0.033), PE 34:2 (adj. p=0.033) and PC 34:4 (adj. p=0.048). Five additional metabolites were significantly elevated in ASD_CASK_ (Table S8G) including PE e 38:4 (adj. p=0.012), PE 38:3 (adj. p=0.0010) and PE 36:4 (adj. p=0.0015).

## DISCUSSION

As the identification of both genetic and environmental factors in different disease conditions such as ASD and other NDDs is increasing, there is a need for efficient, multiplexable platforms to model their effects and interactions during differentiation of neural progenitors. Here, we describe our study using FFED coupled with RNA-seq to identify affected biological pathways in a human iPSC derived model of early neurodevelopment, followed by a layered analytical pipeline. We find modest but significant changes at the pathway level after low-grade exposure with six selected environmental factors and showed the feasibility of separating the effects at different levels of complexity, vis-à-vis global, clinical background, and cell line effects, independently and as two-way interactions.

A means to highlight the efficiency and utility of the FFED would be to compare the input sample count in similar transcriptomic studies. While we performed RNA-seq on 32 samples and in-depth analysis to study the effects of six environmental factors, a theoretical full design study would require 120 samples to study 6 factors in 4 cell lines using 5 technical replicates. Similarly, a conventional experimental design to investigate the effects of Pb on human neural progenitors at two different concentrations and multiple time-points used 60 samples (Jiang et al., 2017). A recent report analysing the effects of a mixture of endocrine disrupting chemicals on neurodevelopment and language delay in human cerebral organoids and *in-vivo* models, generated vast insights on perturbed regulatory networks (Caporale et al., 2022). The FFED, would not only make it possible to analyse such interaction effects, but simultaneously decipher independent effects too, creating an in-depth transcriptomic profile with the same or even lower levels of sampling. As in the future, we need to better model complex mixtures of environmental factors across different doses and time points *in-vitro* (Austin, 2019; Caporale et al., 2022; Curtin et al., 2018), the FFED approach will prove advantageous for such research.

In addition to showing the usefulness of the FFED method to increase the experimental efficiency for transcriptomics or other high-throughput techniques, we have also gained insight into the biological mechanisms of the studied environmental factors. Several of the pathways identified to be changed have been linked to ASD through genetic studies (Eyring and Geschwind, 2021; Satterstrom et al., 2020). These pathways were found to be differentially modulated by the six environmental factors, however, such effects were predominantly driven by the exposures of Pb and FH. It is important to note here that these effects represent sustained, steady state transcriptomic responses following low-grade environmental exposure for 5 days and therefore provide stable insights with greater translational value.

We detect several relevant transcriptomic changes in the neural progenitors after low-grade exposure to Pb, primarily related to synaptic function. Our findings support previous reports on Pb exposure effects, a well-known neurotoxicant linked to cognitive impairments, ASD, and other NDDs (Heyer and Meredith, 2017; Neal and Guilarte, 2010). *In-vitro* studies looking at the mechanism of Pb action with the help of primary neuronal cultures showed that Pb can affect synaptic plasticity (Neal and Guilarte, 2010) and the expression of multiple ASD susceptibility genes (Jiang et al., 2017). These reports suggest that Pb induces modulation of selected genes implicated in vesicle release, neurite growth, synaptogenesis, and transcription factors involved in neuronal differentiation (Sanchez-Martin et al., 2013). Our analyses align with the earlier studies, as we detect upregulated pathways related to axonogenesis and synaptogenesis. Interestingly, we also observed a large cluster of downregulated pathways related to DNA repair, cell cycle/division, and nucleosome remodelling. While such effects are known cytotoxic changes induced by Pb exposure (Gidlow, 2015; Hartwig, 1994; Jannuzzi and Alpertunga, 2016; Senut et al., 2012), these changes have also been previously identified in ASD (Courchesne et al., 2019; Markkanen et al., 2016; Mossink et al., 2021; Satterstrom et al., 2020).

Another exposure resulting in large pathway level changes was FH, a commonly prescribed SSRI for depression and anxiety disorders. We show that FH modulates transcriptomic pathways related to lipid metabolism, which we also functionally validated using ESI-MS. We indicate an increase in lipid-based metabolites following FH exposure in differentiating neural progenitors. The importance of lipids in the human brain is well known, which has the highest lipid content after adipose tissue (Hamilton et al., 2007). Neural progenitors are rich in lipid droplets and their abundance influences the proliferative state of such progenitors, which may also contribute to cell fate determination (Ramosaj et al., 2021). In neural embryonic stem cells, FH exposure at different time points of neural stem cell differentiation was shown to alter cell type markers (de Leeuw et al., 2020). Mechanistically, FH binds to the transmembrane domain of the dimerised neurotropic tyrosine kinase receptor 2 (TKRB), otherwise responsible for long-term potentiation thorough brain derived neurotrophic factor (BDNF) signalling, which we also detect as altered after FH exposure (Casarotto et al., 2021).

How the changes in lipid metabolism during differentiation of neural progenitors would affect outcomes such as ASD later in life, requires further investigation. Nevertheless, it is still intriguing to note that deviations in lipid profiles have been previously observed in individuals with ASD. In a systematic review by Esposito and colleagues, 37 studies linking ASD to abnormal lipid profiles were identified (Esposito et al., 2021), showing an association between ASD and hypocholesterolaemia but unclear association for fatty acids. Furthermore, recently, hypolipidemia was identified in 367 individuals with an ASD diagnosis (Tierney et al., 2021). We also noted that the baseline concentrations for many of the affected lipids were lower in the Non-ASD cell line, and these get elevated following short-term FH exposure in both groups. While transcriptomic enrichment of lipid-based pathways has been noted in ASD (David et al., 2016) with an increasing interest in defining an ASD subtype based on dyslipidaemia (Luo et al., 2020), we provide evidence that FH exposure in early development could alter metabolomic pathways, thereby suggesting the investigation of long-term exposure effects as well as comparisons with other SSRIs should be made.

Interestingly, evidence of FH effects on lipid metabolism has started gaining traction. For instance, long term exposure to FH for 2 years in male rhesus macaques decreased polyunsaturated fatty acids in the medial prefrontal cortex (Tkachev et al., 2021). An increase in serum lipid levels was observed in patients of clinical depression following treatment with FH for 8 weeks (Pan et al., 2018). Even though there has been uncertainty regarding the direction of such changes, a report on FH effects in a murine model of depression revealed that the nature of the biological sample used in a metabolomic investigation is a key determinant of the observed lipid patterns (Zhao et al., 2019).

We also provide proof that when investigating effects of environmental exposures using *in-vitro* cellular models, diverse genetic backgrounds need to be taken into consideration. When analysing the effects separately for the clinical background and cell lines, several significantly enriched pathways were found to be predominantly affected by the exposure of Pb and FH, with both independent and combined effects. Remarkably, with a higher network diameter, Pb had a larger upregulatory response in the ASD group (ASD_CASK_, ASD_HNRNPU_), while FH had such an impact on the Non-ASD group (CTRL_Male_, CTRL_Female_). This could signify that the ASD group is more vulnerable to Pb, and the Non-ASD group is to FH, thereby showing a larger transcriptomic response to the factors respectively. Additionally, few pathways modulated by VPA, BPA, EtOH and Zn-were also detected. What was particularly interesting was the ability of BPA to induce differential exon usage events in differentiating human neural progenitors. To find such changes by an already established endocrine disruptor (Caporale et al., 2022) with previously reported effects on alternative splicing (Kim et al., 2021), emphasises the necessity to carry out further research in the context of NDDs.

Our study shows that the FFED is a promising design to generate multiplexed data, when evaluating the effects of several factors on biological outcomes. We have shown that it is possible to translate from exploratory analysis to functional validation with the FFED. We saw robust effects after the Pb and FH exposures in the four cell lines included in the study, but only modest pathway level changes for the other four exposures. A majority of such pathway level changes were also unique to one cell line, which demonstrated a need for larger studies with several cell lines, selected based on genetic and clinical profiles. In conclusion, we have provided evidence for an efficient and multiplexable resource that can be used for better understanding the functional ramifications of environmental factors and gene-environment interactions in ASD and other clinical conditions with neurodevelopmental links. A molecular level understanding of the biological underpinnings of both genetic and environmental factors is essential for identifying plausible aetiologies and emergent future preventive interventions in ASD.

## SUPPLEMENTAL INFORMATION

Supplemental information can be found online.

## ACKNOWLEDGMENTS

The authors are grateful to the donors of the cell lines and for their participation in this research. The authors acknowledge support from the National Genomics Infrastructure in Stockholm funded by Science for Life Laboratory, the Knut and Alice Wallenberg Foundation and the Swedish Research Council, the SNIC/Uppsala Multidisciplinary Center for Advanced Computational Science for assistance with massively parallel sequencing and access to the UPPMAX computational infrastructure, and the iPS Core facility at Karolinska Institutet for assistance with the generation of iPSC and NES cells. The project was supported by the Swedish Research Council (A.F., and K.T.), Swedish Foundation for Strategic Research (A.F., and K.T.), the Swedish Brain Foundation – Hjärnfonden (A.F., and K.T.), the Harald and Greta Jeanssons Foundations (K.T.), Åke Wiberg Foundation (K.T.), Strategic Research Area Neuroscience Stratneuro (K.T.), The Swedish Foundation for International Cooperation in Research and Higher Education STINT (K.T.), and Board of Research at Karolinska Institutet (K.T.). Open access funding provided by Karolinska Institutet.

## AUTHOR CONTRIBUTIONS

Conceptualization: A.A., M.B., and K.T.; Methodology: A.A., M.B., and K.T.; Software development: A.A., M.B., and K.T.; Validation: A.A., C.M., I.L. and K.T.; Formal Analysis: A.A., M.B, C.M., M.O., D.L, M.W. and F.M., ; Investigation: A.A., M.B. and C.M.; Resources: I.L. and K.T.; Data Curation: A.A., M.B., C.M., M.O., D.L., M.W. and F.M.; Writing – Original Draft: A.A. and K.T.; Writing – Review & Editing: A.A. and K.T.; Visualization: A.A., M.O., D.L. and K.T.; Supervision: M.A., A.F., C.O.D., I.L. and K.T.; Project Administration: A.A. and K.T.; Funding Acquisition: K.T.

## METHODS

### RESOURCE TABLE

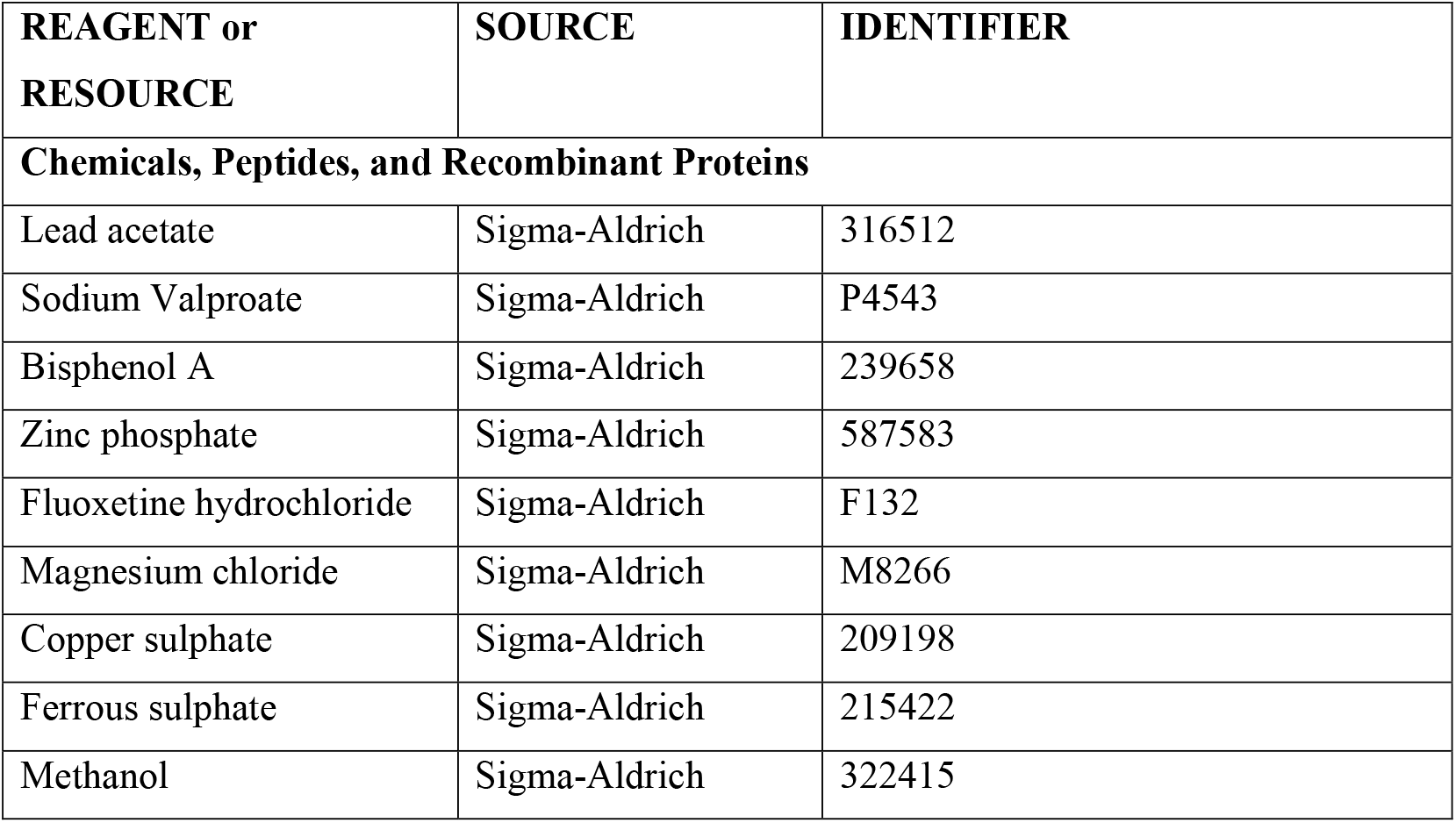
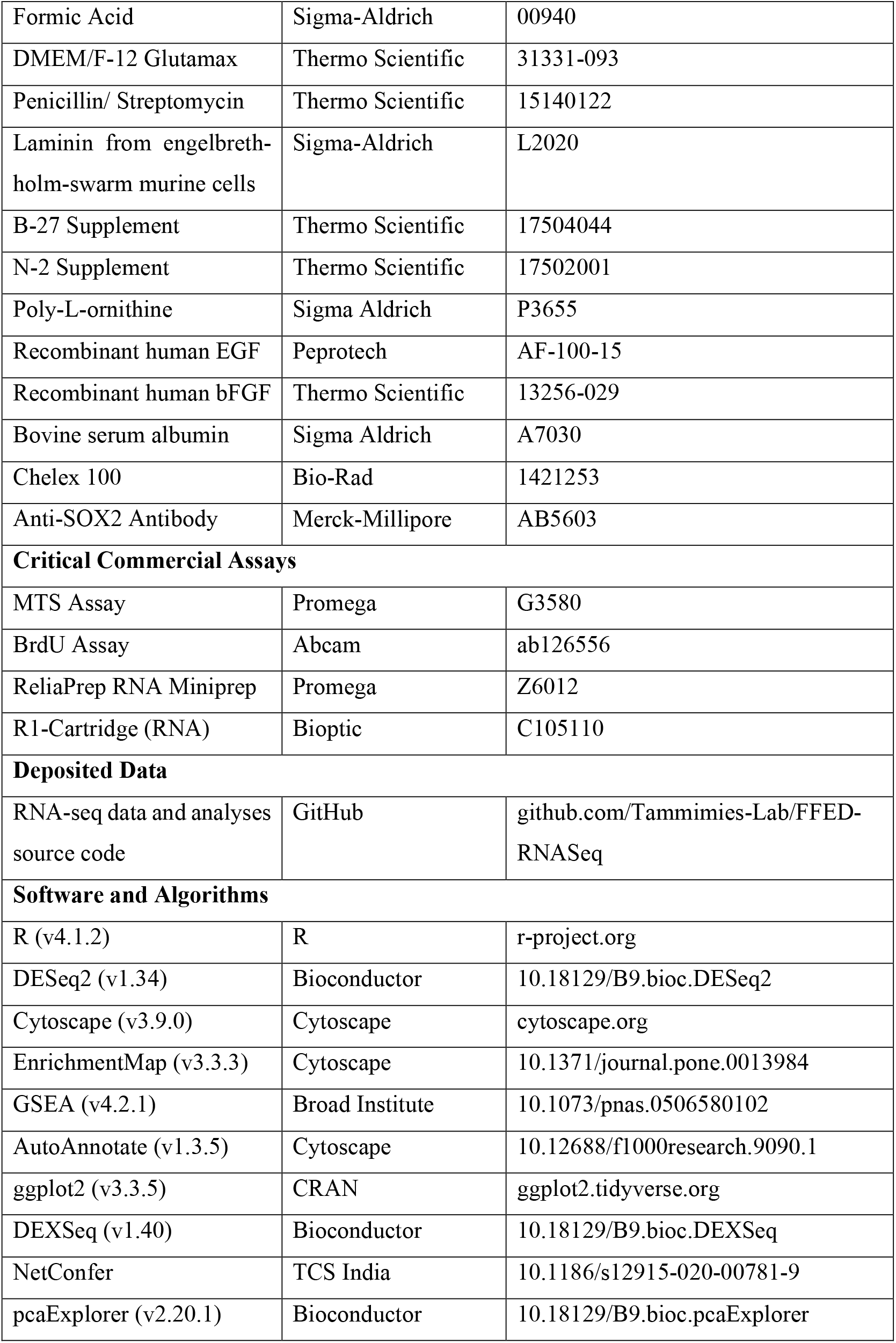
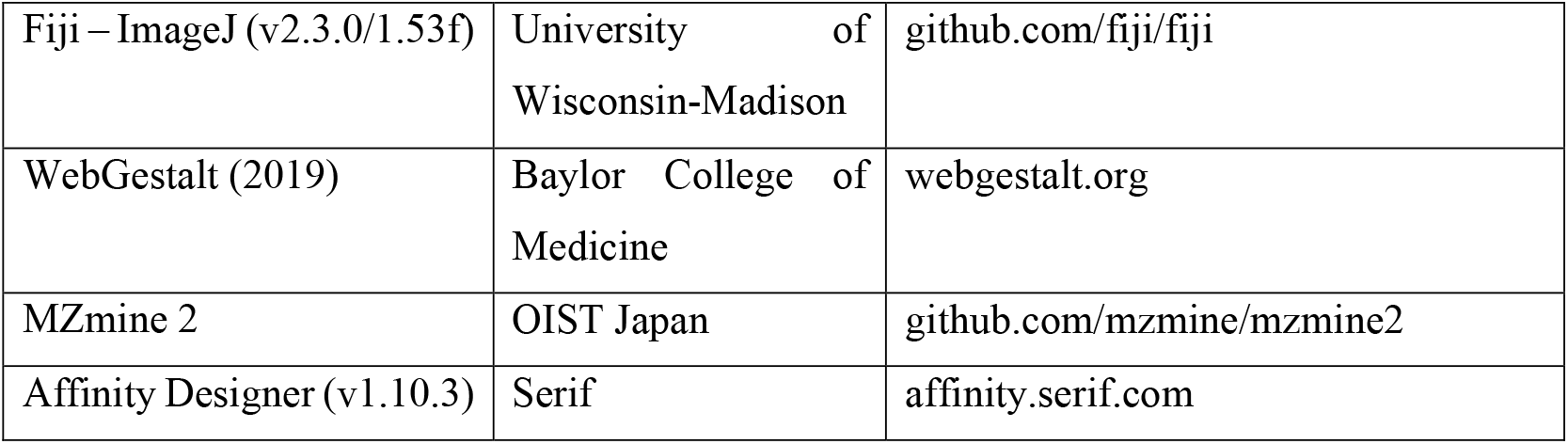

### LEAD CONTACT AND MATERIALS AVAILIBILTY

Further information and requests for resources and reagents should be directed to and will be fulfilled by the Lead Contact, Kristiina Tammimies.

### EXPERIMENTAL MODEL AND SUBJECT DETAILS

#### Cell culture

For the purpose of this study, previously generated and described human iPSC lines from tissue samples of a male: CTRL9II – CTRL_Male_ (Uhlin et al., 2017) and female: AF22 – CTRL_Female_ (Falk et al., 2012) neurotypical donors and two male donors with known genetic variants for NDDs including ASD: ASD12BI – ASD_HNRNPU_ (Mastropasqua et al., 2022) and ASD17AII – ASD_CASK_ (Becker et al., 2020), were used (Figure 1A). Information pertaining to the generation and quality control of the iPSC lines, as well as relevant clinical background are provided in the cited publications. The iPS cells were differentiated into neural progenitor state and maintained as neuroepithelial stem (NES) cells, as previously described (Chambers et al., 2009; Falk et al., 2012). While NES cells are neural progenitors with both neurogenic and gliogenic potential, upon differentiation they are primarily neurogenic unless they are specifically differentiated to glia (Chambers et al., 2009; Falk et al., 2012).

The NES cells were seeded on tissue culture treated plates (Sarstedt), coated with 20 μg/mL poly-ornithine (Sigma-Aldrich) and 1 μg/mL Laminin2020 (Sigma-Aldrich). These were grown in DMEM/F-12 Glutamax basal medium (Gibco) supplemented with 0.05x B27 (Gibco), 1x N-2 (Gibco), 10 U/ml Penicillin-Streptomycin (Gibco), 10 ng/mL recombinant human bFGF (Gibco) and 10 ng/mL recombinant human EGF (Peprotech). Undirected differentiation of NES cells was achieved by the withdrawal of growth factors from the culture medium (Chambers et al., 2009; Falk et al., 2012). The NES cells were seeded on tissue culture treated plates (Sarstedt), coated with 20 μg/mL poly-ornithine (Sigma-Aldrich) and 1 μg/mL Laminin2020 (Sigma-Aldrich). These were grown in DMEM/F-12 Glutamax basal medium (Gibco), that was made zinc free for use in the FFED approach, supplemented with 0.5x B27 (Gibco), 1x N-2 (Gibco) and 10 U/ml Penicillin-Streptomycin (Gibco). When using zinc free basal medium, 1.5 µM zinc sulphate (ZnSO_4_, Sigma) was added to maintain normal zinc concentration in the basal medium (Table S1A). The cells were differentiated for 5 days post induction of differentiation, followed by sampling based on the downstream application. To control for any variance introduced by experimentation, the experiments were performed by the same cell culture scientist in a single experimental batch with passage matched NES cells and all samples were collected at the timepoint stated in the methods section.

Zinc free DMEM/F-12 Glutamax basal medium was prepared following treatment of 500 mL DMEM/F-12 Glutamax basal medium (Gibco) with 25 g of Chelex 100 (Biorad). Following gentle mixing on an orbital shaker for 1 hour at room temperature, the pH of the medium was adjusted to 7.0 – 7.4 using hydrochloric acid (HCl, Sigma) and sterile filtered using a 0.02 µ filtration unit (Sarstedt). As Chelex 100 is not a specific chelator for zinc ions but rather chelates all positively charged metal ions, other metal ions had to be restored in the medium by addition of a self-formulated metal supplement. To prepare 1 mL of metal supplement for 500 mL of medium, add 525.2 µL of 1 M calcium chloride (CaCl_2_, Sigma), 0.65 µL of 1 mg/mL copper(II) sulphate (CuSO_4_, Sigma), 4.17 µL of 100 mg/mL iron(II) sulphate (FeSO_4_, Sigma) and 301.5 µL of 1 M magnesium chloride (MgCl2, Sigma) to 168.5 µL of Milli-Q water. The metal supplement was sterile filtered using 0.02 µ syringe filter prior to thoroughly mixing with the basal medium.

## METHOD DETAILS

### Fractional factorial experimental design (FFED)

Based on our aim to study six exposures with two levels each (untreated control and treated), we selected the L8 Orthogonal Array based FFED (Mee, 2009). The cell culture media was spiked with the environmental factors as per the FFED and determined treatment concentration, to mimic sustained low-grade exposures of the environmental factors *in-vitro* (Table S1A).

### Cytotoxicity assays for environmental factors

We evaluated the cytotoxicity of lead (Pb), valproic acid (VPA), bisphenol A (BPA), ethanol (EtOH), fluoxetine hydrochloride (FH) and zinc deficiency (Zn-) in our cell lines. The treatment concentration ranges (0.00 – 30.00 µM or 0.00 – 200.00 mM) for the selected environmental factors were determined based on those previously reported in different cell types and tissue samples from the literature. (Table S1), followed by testing in our cell lines using the MTS cytotoxicity assay (Promega).

The NES cells were exposed to a concentration range of the environmental factor being studied. Growth factors were withdrawn from the cell culture, to initiate differentiation. The exposure media was changed every second day. At 24- and 120-hours (5 days) post exposure, the cells were treated with MTS (3-(4,5-dimethylthiazol-2-yl)-5-(3-carboxymethoxyphenyl)-2-(4-sulfophenyl)-2H-tetrazolium) reagent, based on instructions from the manufacturer. After 3 hours of incubation at 37°C and 5% CO_2_, absorbance was recorded at 490 nm using a spectrophotometric microplate reader. The same experimental approach was repeated when checking the selected concentrations for the environmental factors using the FFED (Table S1A).

### Immunocytochemistry

The stem cell specific marker, SOX2 was visualised using immunocytochemistry. Cells were cultured on glass coverslips with exposure to the environmental factors in the FFED for 120 hours (5 days) and fixed for 20 minutes in 4% paraformaldehyde. The primary antibody used was SOX2-AB5603, 1:1000 (Merck-Millipore). All images were taken with LSM 700 Zeiss Confocal Microscope (Zeiss Plan-Apochromat 63×/1.40na Oil DIC Objective M27), with 63× magnification at 1024-× 1024-pixel (pxl) resolution, resulting in an aspect ratio of 0.099233 μm per pixel. Fiji – ImageJ (v2.3.0/1.53f) (Schindelin et al., 2012) was used to estimate SOX2 positivity, normalised against total cellular nuclei. Quantification of percentage positivity was done in R (v4.1.2) (R Core Team, 2020).

### Cell proliferation

To analyse differences in cellular proliferation, when exposed to the environmental factors in the FFED, the BrdU assay (Abcam) was performed at 24 hours and 120 hours (5 days) post induction of differentiation. 24 hours prior to the read-out, 1X-BrdU reagent was added to the cell culture vessels for incorporation by incubation at 37°C and 5% CO_2_. For the read-out, the culture media was aspirated, and cells were fixed with the supplied fixing solution. This was followed by exposure to the anti-BrdU antibody (primary antibody), followed by incubation at room temperature for 1 hour and washing with the supplied plate wash buffer. The cells were incubated with the HRP-tagged secondary antibody at room temperature for 30 minutes followed by TMB exposure and recording absorbance at 450 nm. Quantification of percentage proliferation was done in R (v4.1.2) (R Core Team, 2020).

### Bulk RNA sequencing (RNA-seq)

Cell lysates were collected on day 5 of exposure (N=43). At least one biological replicate was included for each of the four cell lines (n=32). An additional full biological replicate was collected for CTRL_Male_ (n=8) and a partial for CTRL_Female_ (n=4) to account for cell culture related effects. RNA samples were extracted using a spin-column based kit (Promega) and following manufacturer’s instructions. RNA integrity was determined using a fluorescence based micro-capillary detection system (Qsep100, Bioptic) using an RNA specific kit (Bioptic) and following manufacturer’s guidelines. RNA samples with an RNA quality number (RQN) greater than 9 were selected for library preparation and sequencing.

Library preparation and sequencing were performed at the National Genomic Infrastructure (NGI), Stockholm. The sequencing library was prepared using the Illumina TruSeq RNA RiboZero GOLD kit. Pooling and sequencing were done on NovaSeq6000 (NovaSeq Control Software 1.6.0/RTA v3.4.4) with a 2×151 setup using ‘NovaSeqXp’ workflow in ‘S4’ mode flowcell. The Bcl to FastQ conversion was performed using bcl2fastq (v2.20.0.422) from the CASAVA software suite. Results from the best practice bioinformatics nf-core/RNAseq pipeline of the NGI, were used for further analysis (Ewels et al., 2020). In short, quality control of read sequences was performed with FastQC, followed by preparation for alignment using UMI-tools (extraction of unique molecular identifiers), Trim Galore! (adapter and quality trimming), BBSplit (removal of genomic contaminants) and SortMeRNA (removal of ribosomal RNA). Read alignment was completed using STAR aligner. Differential gene expression (DGE) analysis was performed using the DESeq2 package (v3.14) (Love et al., 2014) in R (v4.1.2) (R Core Team, 2020).

The analysis for the detection of differentially expressed genes (DEGs) was stratified depending on several levels of complexity. In the *Level I – Global Effects*, comparisons across all cell lines, independent of genetic background were made and DEGs and globally effected pathways were identified. *Level II – Clinical Background Effects*, comparisons based on the clinical background of the iPSC lines were made to identify DEGs. Here, the neurotypical male and female iPSC lines belonged to the non-ASD group, while the male ASD iPSC lines belong to the ASD group. At *Level III – Cell Line* Effects, comparisons were made between the control and treatment groups of each individual cell line included in the study. Here, the linear model used was *design = ∼ exposure*. The data was filtered based on the level of analysis, prior to the application of the linear model. Lastly, *Level IV – Interaction Effects*, were analysed where comparisons were made across all cell lines, both independent as well as dependent on clinical background, for the three possible interactions as per the FFED. This included interactions between Pb and Zn, VPA and FH, and BPA and EtOH. The linear model used was *design = ∼ cell line + replicate + interaction + cell line*interaction*. Based on instructions provided by the package developer, the PCA was performed and were exported for easy visualisation using the pcaExplorer package (v2.20.1) (Marini and Binder, 2019) in R (v4.1.2) (R Core Team, 2020).

### Pathway enrichment

Pathway analysis was done according to previously described protocols (Reimand et al., 2019). Following DGE analysis, RNK files were generated, and Gene-Set Enrichment Analysis (GSEA, v4.2.1) (Subramanian et al., 2005) was performed for biological processes, molecular function, and cellular components, with a significance threshold of FDR<0.05. The scored gene ontology information (biological processes, MSigDB v7.4) was imported into Cytoscape (v3.9.0) (Shannon et al., 2003) and visualised using the EnrichmentMap (v3.3.3) (Merico et al., 2010) app. The generated clusters with FDR adjusted p values (q values) were subsequently labelled using the AutoAnnotate (v1.3.5) (Kucera et al., 2016) app, to identify enriched pathways, that were both differentially upregulated and downregulated. Summary networks were also created following the annotation. The generated network(s) were exported as PDFs and aesthetically organised using Affinity Designer (v1.10.4).

Network parameters for the generated gene ontology biological processes (GOBP) networks were analysed using the online tool NetConfer (Nagpal et al., 2020). With the “WF1: Assess similarity of network components” workflow, lists were generated to indicate unique and shared GOBP terms across the compared networks. The “WF2: Identify and compare key nodes” workflow was used to compute global network properties including total nodes, total edges, diameter, density, clustering coefficient and average degree.

### Gene enrichment

We used the publicly available BrainSpan (Miller et al., 2014) RNA-sequencing dataset that contains spatio-temporal gene expression data pertaining to typical neurodevelopment to analyse the enrichment of DEGs in certain time and regions. After removing low quality samples (RIN < 9.0) from the dataset, and genes with low expression (< 1 FPKM in at least 2 samples) and with low variable expression between samples (> 0 FPKM less than 50% of samples and coefficient of variance < 0.25) (Eising et al., 2019), a total of 344 samples and 24434 genes were screened for further analysis. Four brain regions and eight developmental periods from 8 PCW (weeks post conception) to 40 years of age, were defined (Lin et al., 2015). Hierarchical clustering was performed to group significant gene lists to 2 clusters separately, based on their standardized mean expression pattern across all brain regions and timepoints. To identify if there were any significantly enriched gene clusters in any brain regions or time periods, the mean expression of random genes with similar numbers of selected cluster at each region and time period were calculated. The p value was based on the proportion of selected gene clusters’ mean expression that were higher than the random genes’ mean expression after 10,000 times of permutation. FDR (Benjamini and Hochberg, 1995) was used to adjust p values for multiple comparisons and were reported.

To examine for enrichment of the significant DEGs following exposure to the environment risk factors in publicly available ASD and NDD related gene lists, genetic data was obtained from SFARI gene (v2021 Q3) (Banerjee-Basu and Packer, 2010), Genetic epilepsy syndromes (EPI) (v2.489) panel (Stark et al., 2021), Intellectual disability (v3.1500) (ID) panel (Stark et al., 2021) and a developmental gene list (Li et al., 2021). Enrichment analysis based on hypergeometric testing was performed using the phyper function in R (R Core Team, 2020). The background gene list for the analysis was generated from the RNA-seq expression data, for expressed genes with a base mean>20.

### Differential exon usage

Differential exon usage (DEU) analyses were performed using DEXSeq package (v1.36.0) (Anders et al., 2012). Flattened annotation file was created using provided python script excluding the aggregate exon bins and exon counts were calculated using provided python script. For the global effects (*Level I*) analysis, the cell line factor was used as a blocking factor using the linear models *formulaFullModel = ∼ sample + exon + cell line:exon + exposure:exon and formulaReducedModel = ∼ sample + exon + cell line:exon*. Therefore, DEU exon bins were called based on differences after treatments and differences between cell line gene expression backgrounds was blocked from the analysis. The analysis was repeated for ASD (ASD_CASK_, ASD_HNRNPU_) and non-ASD cell lines (CTRL_Male_, CTRL_Female_) again blocking gene expression backgrounds between cell lines. Finally, the analysis was performed for all the cell lines separately per treatment by *∼ sample + exon + exposure:exon*. A gene was called to have evidence for DEU, if it had at least one exon bin differentially used between conditions. The difference was considered significant with FDR (Benjamini and Hochberg, 1995) adjusted p<0.05, exon base mean ≥10 and an absolute log fold change ≥1.5. Over-representation analysis (ORA) was performed using the online tool WebGestalt (Liao et al., 2019).

### Direct infusion electrospray ionisation mass spectrometry (ESI-MS)

Metabolomic profiling was performed using the direct infusion probe (DIP) for ESI-MS (Marques et al., 2022) on snap-frozen cell lysates (N=24) at day 5 of exposure to 3.0 µM FH during undirected differentiation of NES cells, with 3 technical replicates each for the treated and untreated groups in all four cell lines: CTRL_Male_, CTRL_Female_, ASD_HNRNPU_ and ASD_CASK_. The cells were stored as pellets in -80 freezer prior to lysing with the electrospray solvent containing 9:1 methanol:water and 0.1% formic acid. To enable comparison, all cell samples had the similar number of cells per volume. The samples were analysed on a QExactive Orbitrap instrument at 140,000 mass resolving power using full scanning between 70 and 1000 Da in untargeted mode. The resulting data was extracted using one minute data per sample and sorted in MZmine2 (Pluskal et al., 2010). Quantification was performed using a one-point calibration and the metabolite concentration in the sample was calculated from the concentration of the corresponding internal standard multiplied with the intensity ratio of the endogenous metabolite and the internal standard.

To test for significance of the resulting metabolite concentrations (µM) following the mass spectrometry analysis pipeline, a linear model was applied to first the complete dataset, then to subset datasets based on the clinical background groups and individual cell lines. Here, the linear model used was *lm(concentration ∼ exposure)*. The data was filtered prior to the application of the linear model, based on the level of analyses: global effects, clinical background effects, and cell line effects. The obtained p values were adjusted for multiple comparisons using the FDR method in R (v4.1.2) (R Core Team, 2020). Significantly modulated metabolites (Untreated vs. Treated) were selected based on a significance threshold of adjusted p<0.05 and were visualised using the ggplot2 package (v3.3.5) (Wickham, 2009) in R (v4.1.2) (R Core Team, 2020).

## QUANTIFICATION AND STATISTICAL ANALYSIS

All statistical analyses were performed in R (v4.1.2). The statistical models and tests used for the analyses are described in the methodology relevant to the experimental technique, in the sections above.

## DATA AND CODE AVAILABILITY

The generated data and utilised code is available on GitHub (https://github.com/Tammimies-Lab/FFED-RNASeq) or available upon reasonable request from the corresponding author.

## FIGURE LEGENDS

## Supplementary Figures

**Figure S1.**
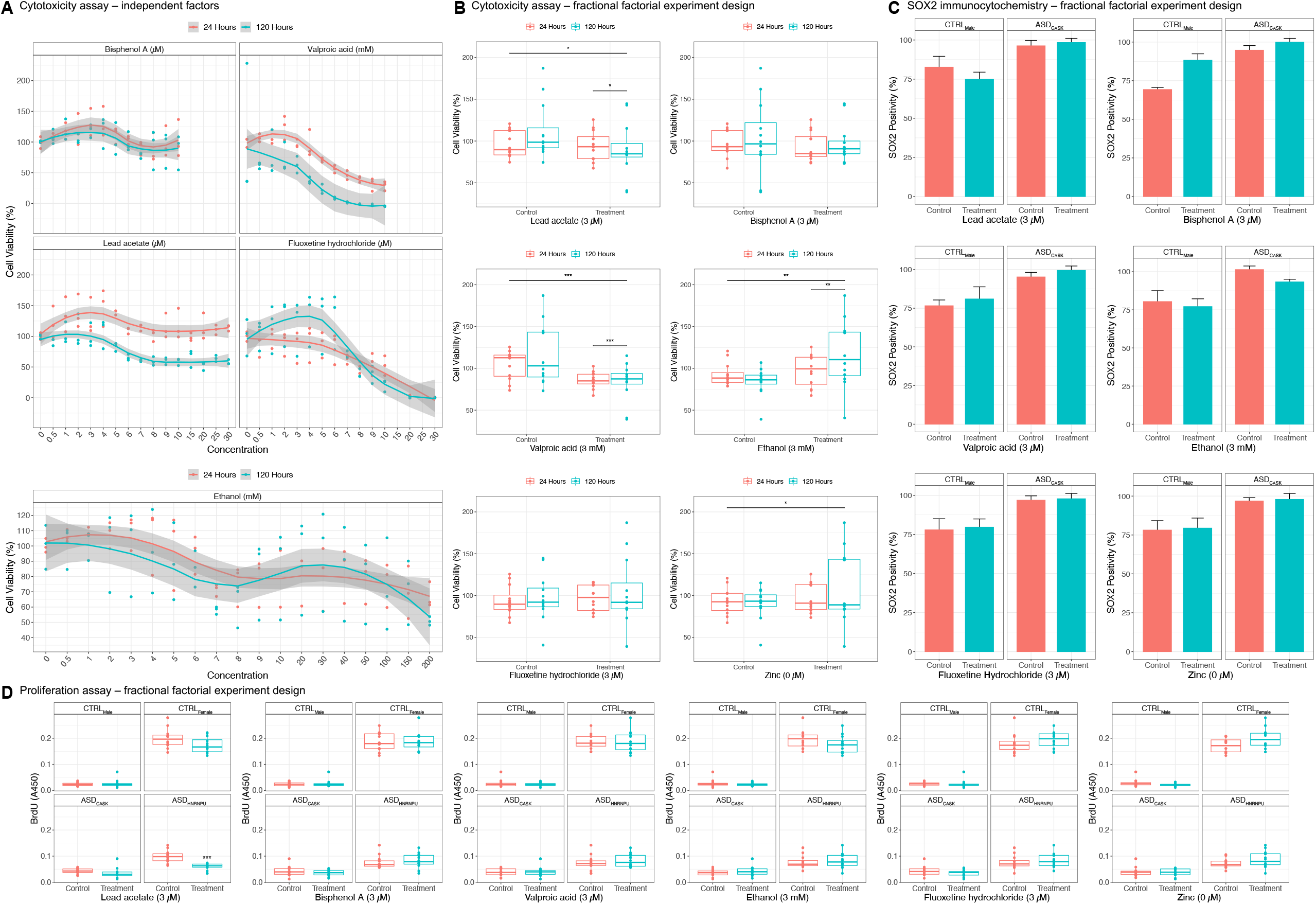
Cellular effects of environmental factors on differentiating neural progenitors. (A) Estimation of cell viability following exposure of lead (Pb), valproic acid (VPA), bisphenol A (BPA), fluoxetine (FH) and ethanol (EtOH) for 24 and 120 hours (5 days) on undirected differentiation of CTRL_Male_ using the MTS cytotoxicity assay. (B) Estimation of cell viability for selected concentrations (from A) of Pb, VPA, BPA, EtOh, FH, and zinc deficiency (Zn-) for 24 and 120 hours (5 days) on undirected differentiation of CTRL_Male_ using the fractional factorial experiment design (FFED) and MTS cytotoxicity assay (Tukey post hoc p *<0.05, **<0.01, ***<0.001). (C) SOX2 positivity following exposures of selected concentrations (from A) of Pb, VPA, BPA, EtOh, FH and Zn-in FFED for 120 hours (5 days) on undirected differentiation of CTRL_Male_ and ASD_CASK_. (D) Changes in cellular proliferation from exposures of selected concentrations (from A) of Pb, VPA, BPA, EtOH, FH, and Zn-in FFED at 120 hours (5 days) on undirected differentiation of CTRL_Male_, CTRL_Female_, ASD_CASK_ and ASD_HNRNPU_ using the BrdU assay (Tukey post hoc p ***<0.001).

**Figure S2.**
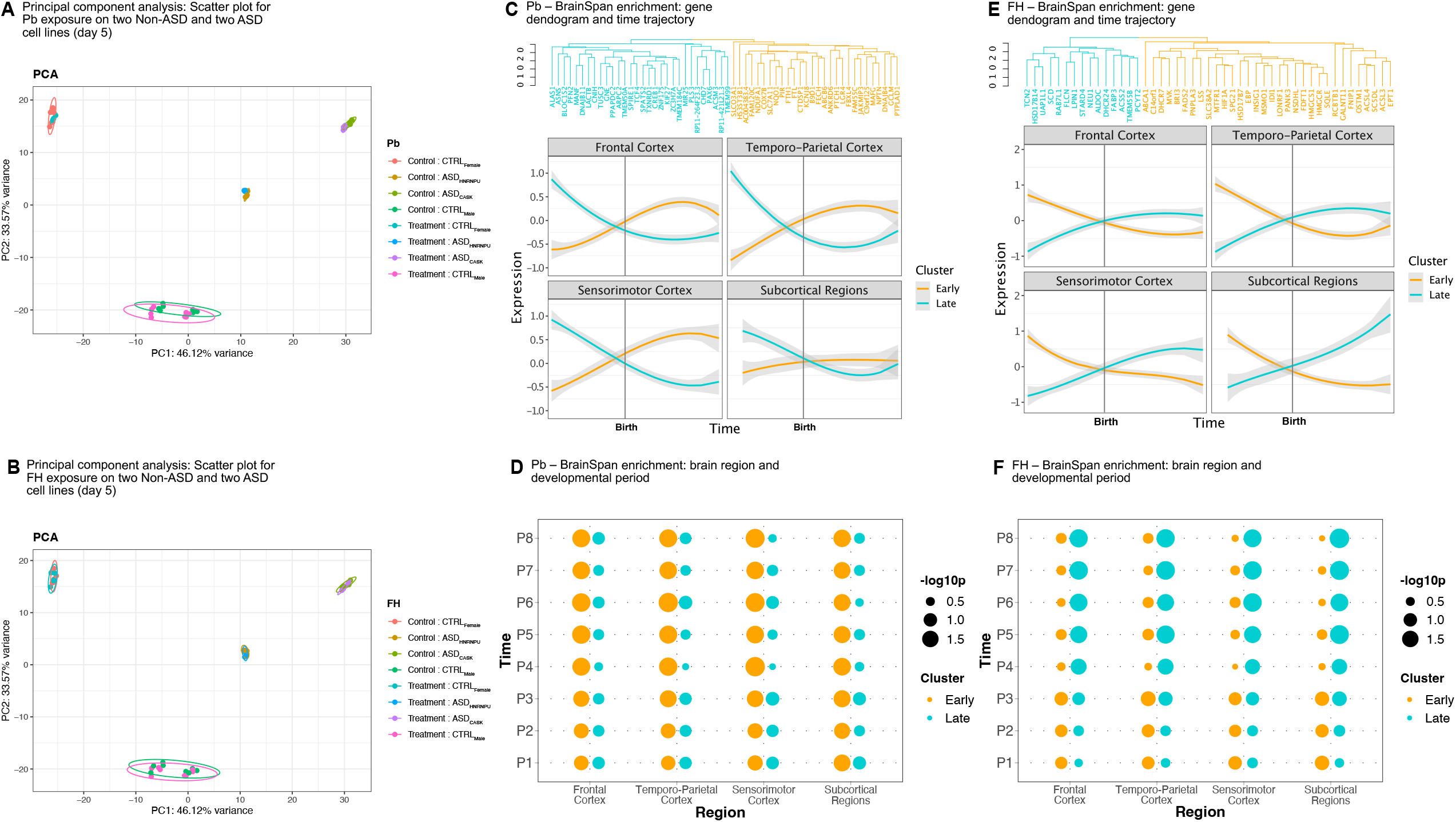
Differential expression and gene enrichment for lead (Pb) and fluoxetine (FH) Principal component analysis (PCA) scatter plot of non-ASD and ASD cell lines exposed to Pb for 5 days. PCA scatter plot of non-ASD and ASD cell lines exposed to FH for 5 days. (A) Cluster dendogram and developmental period trajectory of significant differentially expressed genes (DEGs) following Pb exposure for 5 days, in the Brainspan dataset. (B) Enrichment dot plot of significant DEGs following Pb exposure for 5 days, in the Brainspan dataset across four brain regions (Frontal Cortex, Temporo-parietal Cortex, Sensorimotor Cortex, and Subcortical Regions) for developmental periods (P1: Early foetal<=12 weeks, P2: Early mid-foetal 13-18 weeks, P3: Late mid-foetal 19-24 weeks, P4: Late foetal 25-38 weeks, P5: Infancy 18 months, P6: Childhood 19 months-11 years, P7: Adolescence 12-19 years, P8: Adulthood 20-60+ years). (C) Cluster dendogram and developmental period trajectory of significant DEGs following FH exposure for 5 days, in the Brainspan dataset. (D) Enrichment dot plot of significant DEGs following FH exposure for 5 days, in the Brainspan dataset across four brain regions (Frontal Cortex, Temporo-parietal Cortex, Sensorimotor Cortex, and Subcortical Regions) for developmental periods (P1: Early foetal<=12 weeks, P2: Early mid-foetal 13-18 weeks, P3: Late mid-foetal 19-24 weeks, P4: Late foetal 25-38 weeks, P5: Infancy 18 months, P6: Childhood 19 months-11 years, P7: Adolescence 12-19 years, P8: Adulthood 20-60+ years). (E) Cluster dendogram and developmental period trajectory of significant DEGs following FH exposure for 5 days, in the Brainspan dataset. (F) Enrichment dot plot of significant DEGs following FH exposure for 5 days, in the Brainspan dataset across four brain regions (Frontal Cortex, Temporo-parietal Cortex, Sensorimotor Cortex, and Subcortical Regions) for developmental periods (P1: Early foetal<=12 weeks, P2: Early mid-foetal 13-18 weeks, P3: Late mid-foetal 19-24 weeks, P4: Late foetal 25-38 weeks, P5: Infancy 18 months, P6: Childhood 19 months-11 years, P7: Adolescence 12-19 years, P8: Adulthood 20-60+ years).

**Figure S3.**
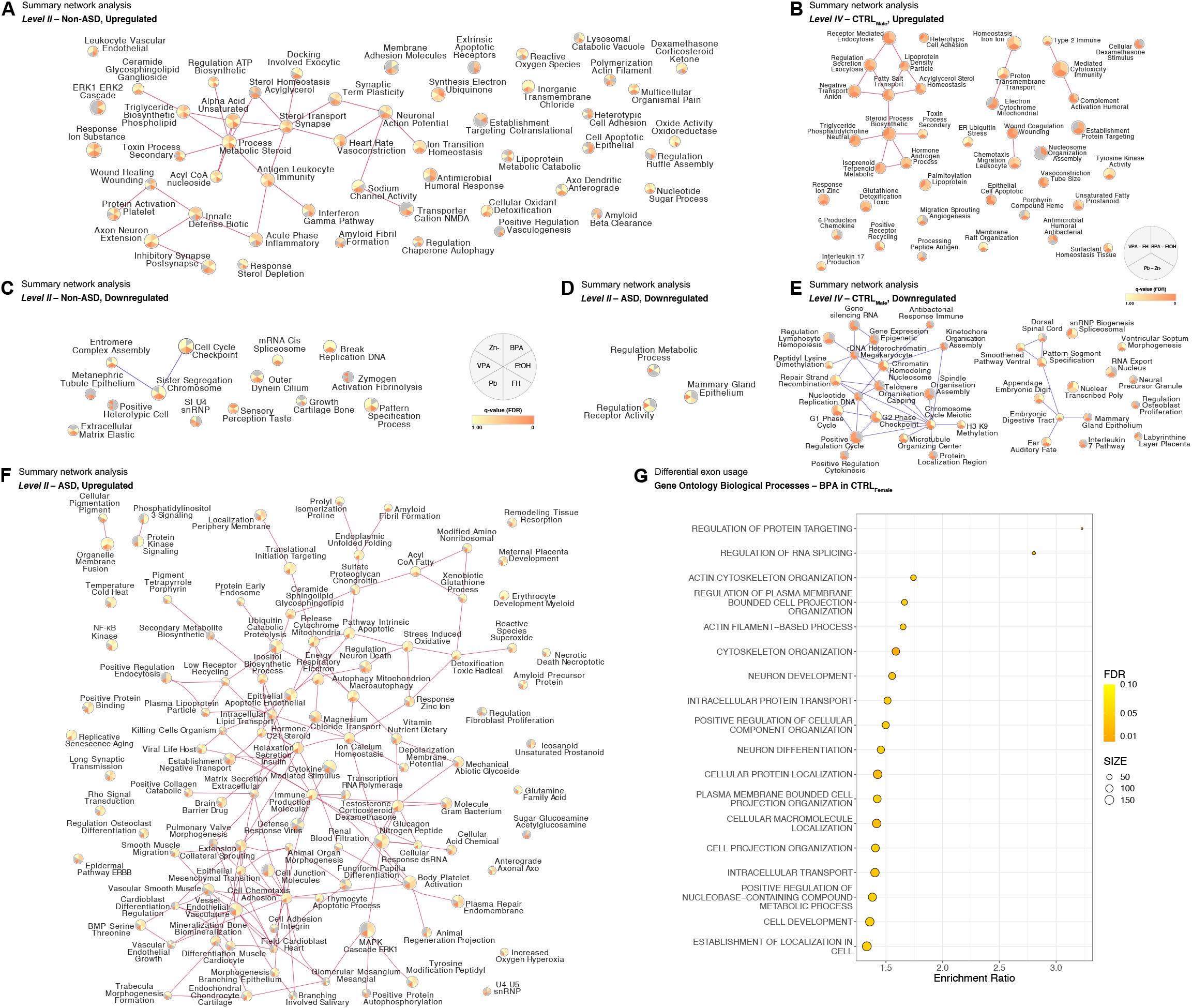
Summary networks and differential exon usage (DEU) of selected environmental factors in differentiating neural progenitors. (A) Upregulated clusters in the non-ASD group (*Level II* analysis). (B) Upregulated clusters for two-way interactions in CTRL_Male_ (*Level IV* analysis). (C) Downregulated clusters in the non-ASD group (*Level II* analysis). (E) Downregulated clusters in the ASD group (*Level II* analysis). (F) Downregulated clusters for two-way interactions in CTRL_Male_ (*Level IV* analysis). (G) Upregulated clusters in the ASD group (*Level II* analysis). (H) Gene ontology biological processes (GOBP) enrichment terms of significant DEU genes (adj. p<0.05) following bisphenol A (BPA) exposure in CTRL_Female_ (*Level III* analysis).

**Figure S4.**
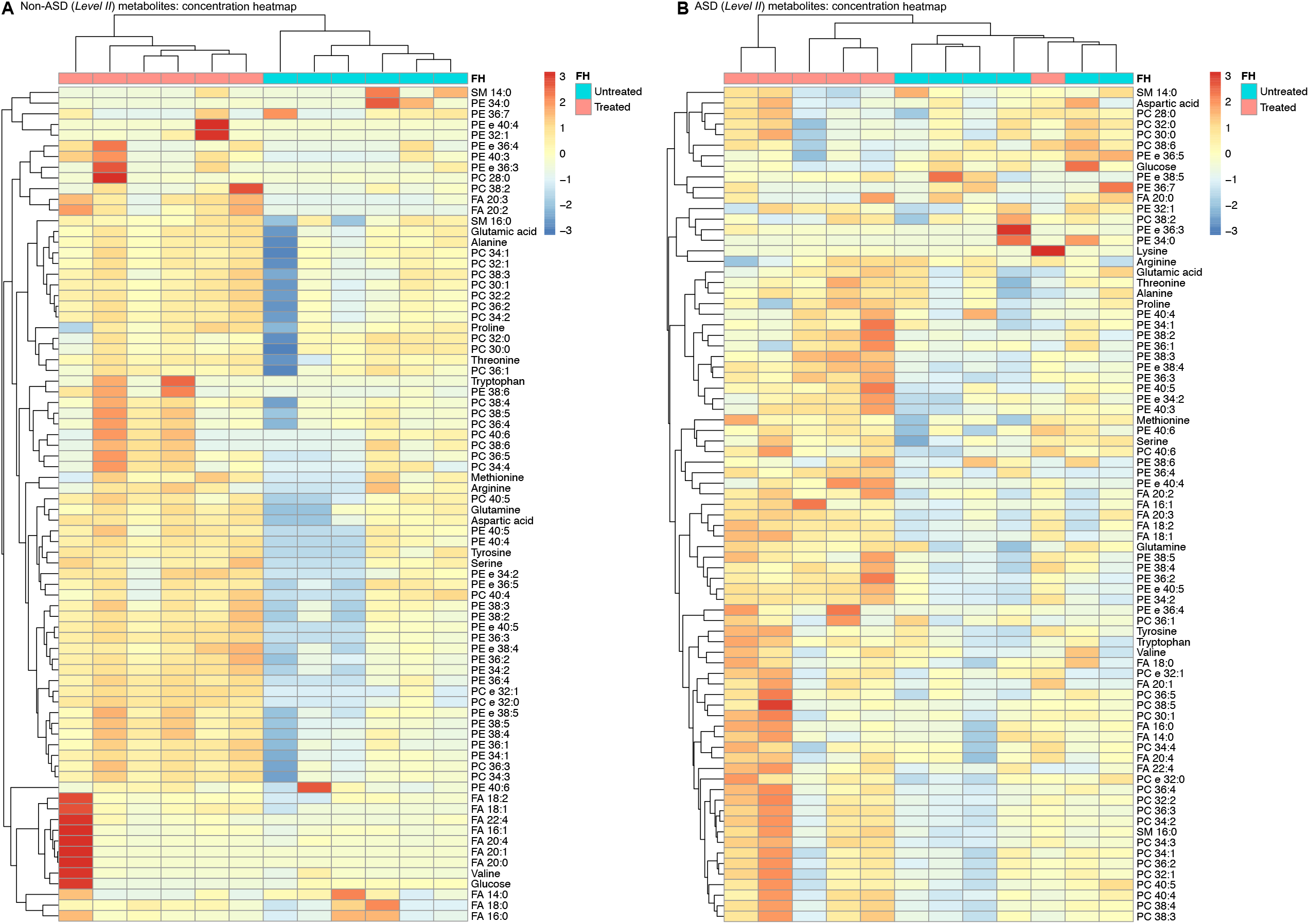
Mass spectrometry-based metabolomics of fluoxetine (FH) exposure. (A) Concentration heatmap of selected metabolites following FH exposure for 5 days in the non-ASD group. (B) Concentration heatmap of selected metabolites following FH exposure for 5 days in the ASD group.

